# O-GlcNAcylation of small heat shock proteins enhances their anti-amyloid chaperone activity

**DOI:** 10.1101/869909

**Authors:** Aaron T. Balana, Paul M. Levine, Somnath Mukherjee, Nichole J. Pedowitz, Stuart P. Moon, Terry T. Takahashi, Christian F. W. Becker, Matthew R. Pratt

**Author notes:** Corresponding Author: Matthew R. Pratt.

## Abstract

A major role for the intracellular posttranslational modification O-GlcNAc appears to be the inhibition of protein aggregation. Most of the previous studies in this area have focused on O-GlcNAcylation of the amyloid-forming proteins themselves. Here, we use synthetic protein chemistry to discover that O-GlcNAc also activates the anti-amyloid activity of certain small heat shock proteins (sHSPs), a potentially more important modification event that can act broadly and substoichiometrically. More specifically, we find that O-GlcNAcylation increases the ability of sHSPs to block the amyloid formation of both α-synuclein and Aβ. Mechanistically, we show that O-GlcNAc near the sHSP IXI-domain prevents its ability to intramolecularly compete with substrate binding. Our results have important implications for neurodegenerative diseases associated with amyloid formation and potentially other areas of sHSP biology.

## Introduction

O-GlcNAcylation (**Figure 1a**), the addition of the monosaccharide *N*-acetylglucosamine (GlcNAc), is an intracellular posttranslational modification (PTM) that occurs on serine and threonine residues (**Figure 1a**) ^1-3^. The installation of this PTM is catalyzed by the enzyme O-GlcNAc transferase (OGT) and it is removed by the glycosidase O-GlcNAcase (OGA)^4^. These enzymes are required for embryonic development in mice and *Drosophila*^5-7^. A variety of proteins have been shown to be O-GlcNAc modified including regulators of transcription and translation, signaling proteins, and metabolic enzymes. The biochemical consequences of most of these modifications are unknown, but limited analyses demonstrated that O-GlcNAc modification can change protein localization, stability, molecular interactions, and activity. One emerging role for O-GlcNAcylation is the inhibition of protein aggregation and the progression of neurodegenerative diseases^8,9^. For example, tissue specific loss of O-GlcNAc through knockout of OGT in neurons results in neurodegeneration^10,11^. Additionally, O-GlcNAcylation levels in the brains of Alzheimer’s disease patients is lower than the brains of age-matched controls. ^12-15^A hallmark of neurodegeneration is the accumulation of misfolded proteins (e.g., tau in Alzheimer’s and α-synuclein in Parkinson’s disease) that form toxic amyloids. Notably, increasing the amounts of O-GlcNAcylation using a small-molecule inhibitor of OGA slows the progressive formation of tau aggregates in mouse models of Alzheimer’s disease^16-19^. O-GlcNAcylation of tau or α-synuclein also directly inhibits the aggregation of these proteins *in vitro*^20-23^. These results suggest that O-GlcNAcylation may play a protective role by directly preventing the aggregation of such proteins and that loss of these modifications may be a factor in the onset of the associated diseases.

**Figure 1.**
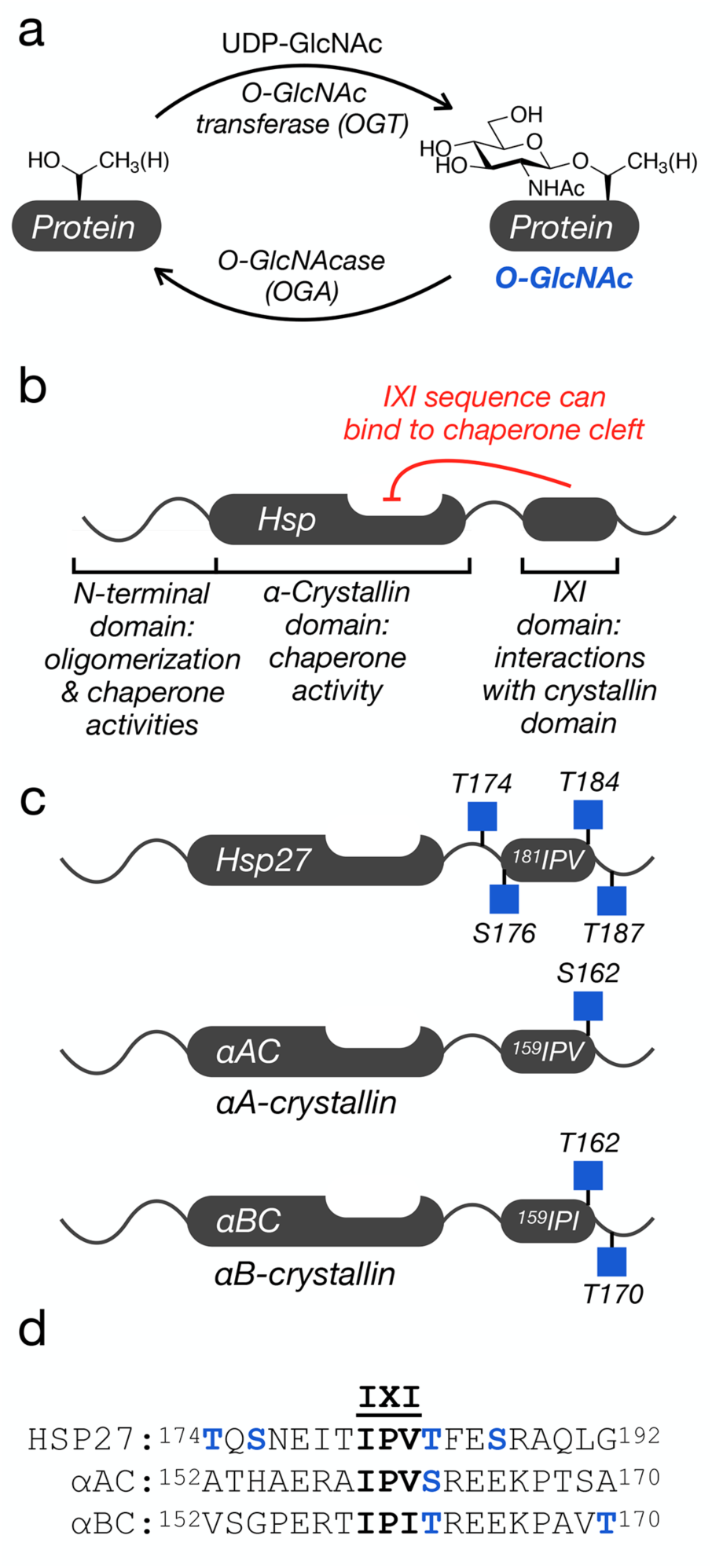
O-GlcNAc modification and the small heat shock proteins (sHSPs). a) O-GlcNAcylation is added to serine and threonine residues of intracellular proteins by O-GlcNAc transferase (OGT) and can be reversed by O-GlcNAcase (OGA). b) Domain structure of a subset of sHSPs, which contains an N-terminal region responsible for protein oligomerization and some chaperone activity, an α-crystallin domain (ACD) that binds to hydrophobic segments, and a C-terminal IXI-domain that regulates the sHSP activity through interactions with the ACD. c) All three sHSPs with a C-terminal IXI domain are O-GlcNAcylated near this sequence in cells and tissues. d) Sequence alignment of the three IXI domains shows a conserved O-GlcNAcylation site (in blue) directly C-terminal to the IXI motif.

Another factor that protects from neurodegeneration is the activity of heat shock proteins, a family of molecular chaperones that are upregulated in response to cellular stress^24,25^. The small heat shock proteins (sHSPs) are a subset that are ATP-independent and function by binding to unfolded/misfolded proteins to prevent their aggregation^26,27^. All sHSPs contain a conserved central α-crystallin domain (ACD) and variable N-terminal and C-terminal domains (Figure 1b) ^28,29^. All of these domains contribute to the formation of large and dynamic protein oligomers that are critical for chaperone activities^30-34^. Of particular interest in neurodegeneration, the ACD domain contains a cleft that appears to be the major site of binding to amyloid-forming proteins, including Aβ, tau, and α-synuclein^30,34-40^. Three human sHSPs - HSP27, αA-crystallin (αAC), and αB-crystallin (αBC) - that have this anti-amyloidogenic activity also contain a tripeptide sequence known as the IXI motif in their C-terminal domains (Figure 1b). The IXI motif can interconvert between an ACD bound form, where the IXI tripeptide is loosely bound to the cleft, and an unstructured form in solution^30,31,40,41^. This IXI-ACD interaction is not required for the formation of large oligomers, but mutants of the IXI motif can have consequences on oligomer stability and dynamics^40,42,43^. Additionally, the contact between the cleft and the IXI-motif can block protein-protein interactions between the sHSP and other proteins, including amyloid substrates^34,40,44^. Therefore, the dynamic regulation of this interaction is proposed to be key contributor to controlling the activity of these sHSPs. Notably, HSP27, αAC, and αBC have been long known to be O-GlcNAcylated^45-48^, and proteomic analysis from cells and tissues has localized endogenous O-GlcNAc modifications to residues very near the IXI domains (Figure 1c) ^45,49-51^. Notably, one of these modification sites, Thr184 in HSP27 and Thr162 αAC/αBC, is conserved between all three proteins (Figure 1d). This led us to hypothesize that O-GlcNAcylation of these sHSPs may inhibit the IXI-ACD interaction and increase their anti-amyloidogenic activity.

Here, we used a combination of synthetic protein chemistry and biochemical analysis to confirm this hypothesis. We first used protein semisynthesis to construct HSP27 bearing individual O-GlcNAc modifications at all four previously identified sites. We then showed that all of these O-GlcNAcylation events improve the chaperone activity of HSP27 against the amyloid aggregation of α-synuclein and that the two modification sites closest to the IXI domain, including the conserved Thr184, displayed the largest increase in this activity. We then applied protein semisynthesis to prepare αAC and αBC with O-GlcNAc at residue 162. Similar to HSP27, O-GlcNAcylation of αAC improved its anti-amyloid activity against α-synuclein, while both unmodified and O-GlcNAcylated αBC completely blocked α-synuclein aggregation. We then tested the possibility that O-GlcNAcylation would also improve chaperone activity against Aβ aggregation and found that all three modified proteins are indeed better chaperones. Because the sHSPs function as oligomers, we also mixed unmodified and O-GlcNAcylated HSP27 at different ratios and found that as little as 25% O-GlcNAcylated monomers is sufficient to induce the increased chaperone activity. Next, we used a variety of biophysical techniques to show that the O-GlcNAcylated IXI domain of HSP27 does indeed display reduced binding to its ACD, providing a mechanism to explain our observations. Taken together, our results demonstrate that O-GlcNAcylation increases the anti-amyloid activity of certain sHSPs and that this protective modification may be lost as a driving event in neurodegenerative diseases. Coupled with our previous work on α-synuclein and the work of others on tau, we believe that O-GlcNAc is a multifaceted inhibitor of amyloid aggregation.

## RESULTS

### Synthesis of O-GlcNAcylated Hsp27

In order to directly test the effect of O-GlcNAcylation at Thr174, Ser176, Thr184, and Ser187 on HSP27 structure and function, we first individually synthesized these modified proteins using expressed protein ligation (EPL) ^52,53^. Traditional EPL relies on a native chemical ligation reaction (NCL) that occurs between a C-terminal thioester and an N-terminal cysteine residue, yielding a native amide bond^54^. HSP27 does not contain a strategically useful cysteine residue close to any of the O-GlcNAc modification sites, so we decided to introduce one at position 173 in the primary sequence, an alanine residue in the native protein. This cysteine mutation allowed us to retrosynthetically deconstruct HSP27 into two fragments, va recombinant protein thioester (**1**) and synthetic peptides (**2**-**6**) containing an N-terminal cysteine residue required for ligation (Supplementary Figure 1). After purification, incubation of **1** with either peptide **2**-**6** in ligation buffer resulted in facile formation of the ligation products. Finally, radical-mediated desulfurization to convert the cysteine required for ligation into the native alanine residue, yielding unmodified and four site-specifically O-GlcNAcylated HSP27 proteins: gT174, gS176, gT184, gS187. HSP27 does contain one native cysteine residue at position 137, which is also converted to alanine in the desulfurization reaction. However, loss of Cys137 has been exploited in the past for semisynthetic access to HSP27^55^, as well as to prevent the formation of covalent HSP27 dimers through a disulfide-bond. This disulfide acts as a regulator of HSP27 activity under different oxidative environments, but is also a complicating factor for the purification and storage of this protein that we wanted to avoid ^56^.

### O-GlcNAcylation improves HSP27 chaperone activity against α-synuclein amyloid formation

As noted above, α-synuclein forms toxic amyloids in Parkinson’s disease, and HSP27 inhibits this process. Therefore, we tested whether O-GlcNAcylation of HSP27 improved the inhibition of α-synuclein aggregation by mixing unmodified HSP27 or one of the O-GlcNAcylated versions (at 1 μM concentration) with α-synuclein (50 μM). We then subjected these protein mixtures to aggregation conditions (agitation at 1,000 rpm, 37 °C) for 7 days. α-Synuclein (50 μM) alone was used as a control. The formation of α-synuclein amyloids was then measured using three different assays. First, we employed the dye thioflavin-T (ThT), which becomes fluorescent in the presence of amyloid fibers (Figure 2a). As expected, unmodified Hsp27 inhibited the aggregation of α-synuclein. Consistent with our hypothesis, all of the O-GlcNAcylated versions of Hsp27 were better aggregation inhibitors, with HSP27(gS176) and HSP27(gT184) having the largest, and statistically significant, effect. Second, we used transmission electron microscopy (TEM) to visualize any aggregates that did form (Figure 2b). We visualized long fibers for α-synuclein alone but smaller amyloid fibers in the presence of unmodified HSP27, and this effect was more pronounced in the presence of HSP27(gT174) or HSP27(gS187). Finally, we found that the best aggregation inhibitors in the ThT assay, HSP27(gS176) and HSP27(gT184), appeared to only form amorphous aggregates. Third, we used proteinase K (PK) digestion to examine the stability of the α-synuclein aggregates. PK displays broad selectivity in the α-synuclein primary sequence and will completely degrade the unfolded protein. However, when amyloids are formed, they inhibit the accessibility of the aggregated region to PK, resulting in stabilized fragments that can be visualized by SDS-PAGE. The resulting banding pattern of the stabilized fragments provides a low resolution picture of the protease-resistant core of the aggregates. Using this assay, we further confirmed that O-GlcNAcylated HSP27 is better at inhibiting the formation of α-synuclein amyloids (Figure 2c). In particular, we discovered that HSP27(gT174) and HSP27(gS187) showed a similar banding pattern as α-synuclein aggregated alone but with reduced intensity of the PK-resistant bands. However, in the presence of HSP27(gS176) or HSP27(gT184), we could only detect very faint bands that were stable to PK, indicating minimal amounts of amyloid formation. Take together, these data show that O-GlcNAcylation results in a site-specific increase in HSP27 chaperone activity.

**Figure 2.**
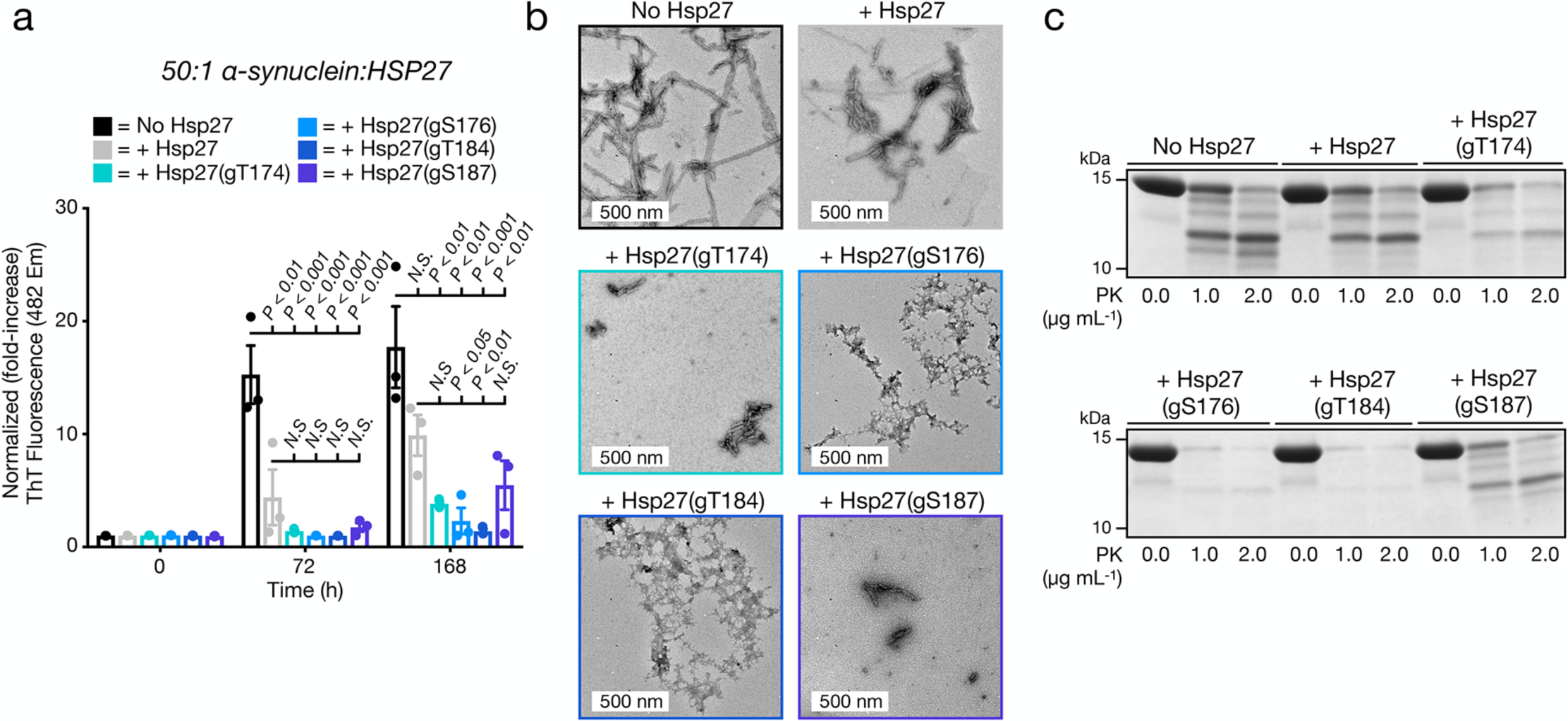
O-GlcNAcylated HSP27 is a better chaperone against α-synuclein amyloid aggregation. a) α-Synuclein alone (50 μM) or in the presence of HSP27 or the indicated O-GlcNAcylated HSP27 proteins (1 μM) was subjected to aggregation conditions (agitation at 37 °C). After different lengths of time, aliquots were removed and analyzed by ThT fluorescence (λ_ex_ = 450 nm, λ_em_ = 482 nm). The y-axis shows fold change in fluorescence compared with α-synuclein alone at t = 0 h. Results are mean ±SEM of three experimental replicates. Statistical significance was determined using a one-way ANOVA test followed by Dunnett’s test (α-synuclein alone or plus HSP27 versus O-GlcNAcylated proteins). b) The same reactions were analyzed by TEM after 168 h. c) The same reactions were subjected to the indicated concentrations of proteinase-K (PK) for 30 min before separation by SDS-PAGE and visualization by Coomassie staining. The persistence of bands correlates with the amount of amyloid formation.

### O-GlcNAcylation is a conserved mechanism for sHSP activation against α-synuclein amyloid formation

Given the increased chaperone activity of HSP27(gT184) and the fact that this O-GlcNAcylated residue is conserved in the IXI domains of αAC and αBC, we next used protein semisynthesis to prepare the analogous O-GlcNAcylated versions of these proteins. Specifically, αAC was retrosynthetically deconstructed into an N-terminal thioester (**7**), residues 1-141, and two peptides (**8** & **9**) (Supplementary Figure 2). Protein **7** was obtained using recombinant expression with the intein technology described above. Peptide thioester **8** was prepared using SPPS on Dawson resin^57^, while glycopeptide **9** was synthesized on Wang resin with a N-terminal selenocysteine residue. We then performed an NCL reaction between peptides **8** and **9**, to yield residues 142-173 of αAC, followed by deprotection of the resulting N-terminal cysteine. Through a subsequent EPL reaction with protein thioester **7**, we obtained the full-length sequence of O-GlcNAcylated αAC. The selenocysteine was then selectively transformed to the native alanine in αAC to yield the O-GlcNAcylated protein with no primary sequence mutations. ^58^ In this case, we required the use of two peptide segments because the recombinant expression of αAC residues 1-156 as an intein fusion resulted in a product that could not be separated by RP-HPLC. Similarly, we prepared αBC from two fragments (Supplementary Figure 3). The first was an intein fusion to residues 1-154 of αBC (**10**), while the second was a synthetic glycopeptide of residues 155-175 (**11**). αBC contains no convenient cysteine nor alanine residues; therefore, we chose to employ γ-thioproline as the cysteine surrogate at the EPL junction^59^. Because αBC does not contain any cysteine residues, we then employed desulfurization to generate glycosylated αBC with no mutations. To obtain the unmodified version of the α-crystallin proteins, we expressed them as full-length N-terminal fusions to an intein and used hydrolysis to remove the intein tag (Supplementary Figures 2 and 3).

**Figure 3.**
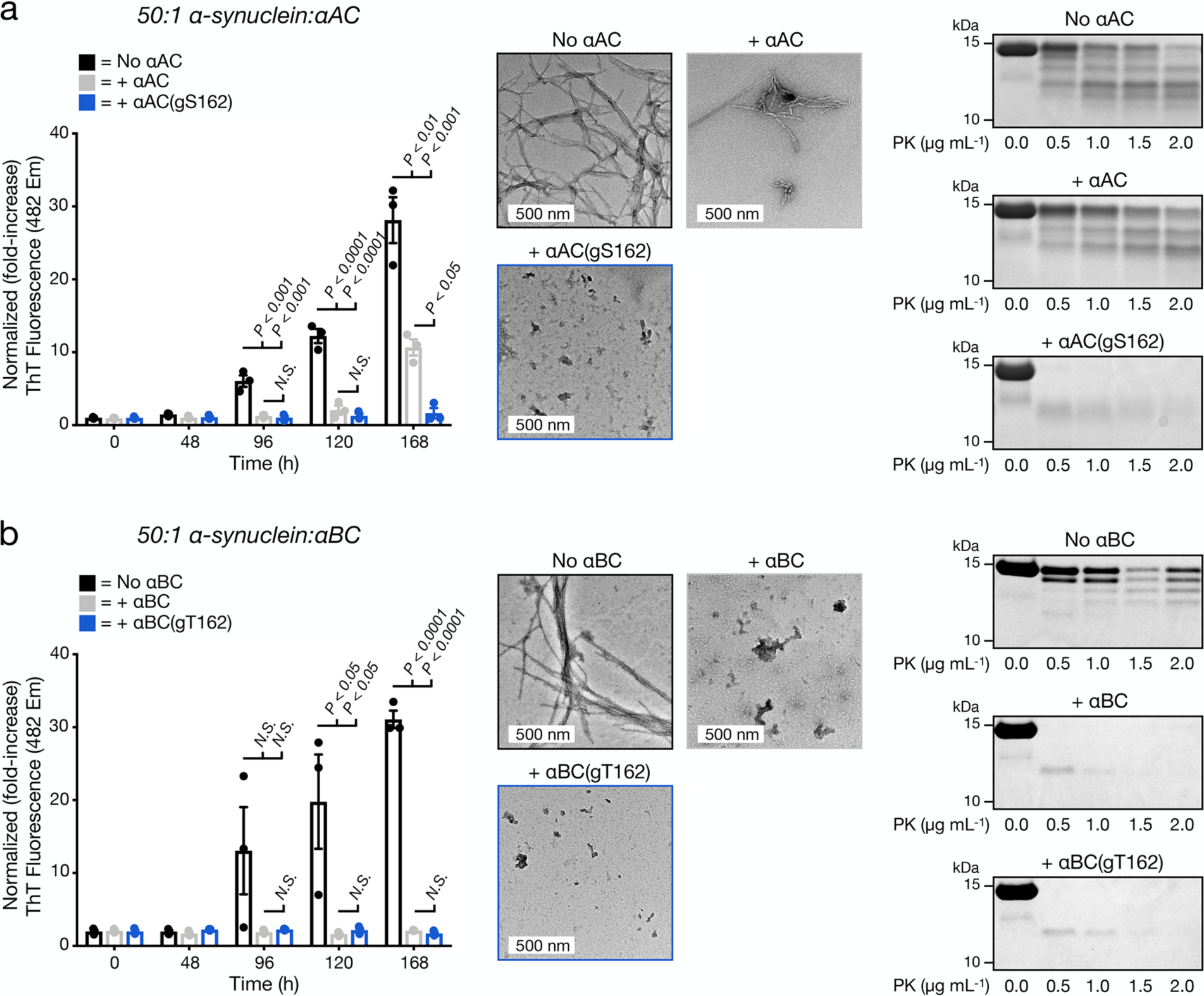
O-GlcNAcylation improves the anti-α-synuclein chaperone activity of αAC/αBC in a protein dependent manner. a) O-GlcNAcylated αAC is a better chaperone against α-synuclein amyloid aggregation. α-Synuclein amyloid formation was measured by ThT fluorescence, TEM imaging, and PK digestion in the absence or presence of αAC or αAC(gS162) as in Figure 2. b) αBC is a strong inhibitor of α-synuclein amyloid aggregation with or without O-GlcNAcylation. α-Synuclein amyloid formation was measured by ThT fluorescence, TEM imaging, and PK digestion in the absence or presence of αBC or αBC(gT162) as in Figure 2.

With these proteins in hand, we next tested the potential for O-GlcNAcylation to increase the chaperone activity of αAC or αBC against α-synuclein amyloid formation. Accordingly, we subjected α-synuclein to the aggregation conditions described above in the absence or presence of either unmodified αAC/αBC or their O-GlcNAcylated variants, αAC(gS162)/αBC(gT162). Again, we performed these aggregation reactions at a 50:1 ratio of α-synuclein to chaperone. Analysis by ThT, TEM, and PK cleavage showed that O-GlcNAcylated αAC was, as predicted, more capable of inhibiting α-synuclein amyloid formation than the unmodified protein (Figure 3a). Notably, by these same measures, both unmodified αBC and the O-GlcNAcylated variant were capable of completely blocking α-synuclein aggregation (Figure 3b), demonstrating that O-GlcNAc at the key conserved Thr residue is minimally capable of activating the chaperone activity of these sHSPS, and is not detrimental to this activity in the case of αBC.

### O-GlcNAcylation activates all three sHSPs against Aβ amyloid formation and can be substoichiometric

sHSPs have also been shown to inhibit the amyloid aggregation of the Aβ(1-42) peptide associated with Alzheimer’s disease, raising the possibility that O-GlcNAcylation of these sHSPs may be a beneficial modification in multiple neurodegenerative diseases. To test this hypothesis, we individually mixed Aβ with either unmodified sHSP or HSP27(gT184), αAC(gS162), or αBC(gT162). In this case, we performed all of these reactions with 10 μM Aβ and 1 μM sHSP. We subjected these mixtures to a ThT plate-reader assay and quantified the onset time of amyloid formation (Figure 4a). As seen in previous publications, Aβ alone formed amyloids very quickly and then precipitated from the reaction solution, resulting in first an increase and subsequent decrease in ThT fluorescence. As expected, we observed a delay in the aggregation of Aβ in the presence of any of the unmodified sHSPs. In the case of the O-GlcNAcylated proteins, we found an even longer onset time for all three sHSPs.

**Figure 4.**
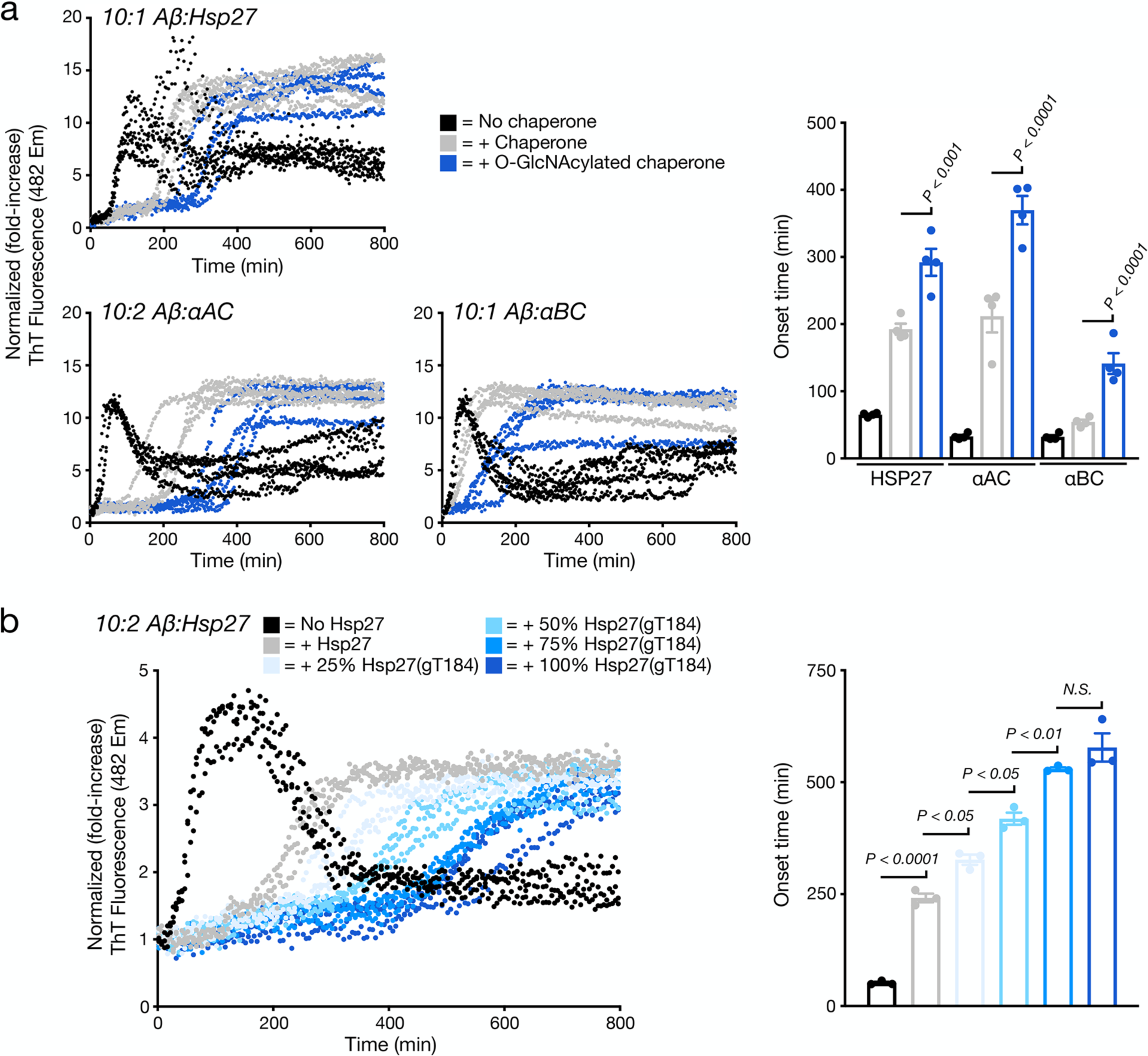
O-GlcNAcylation is a global activator of HSP27, αAC, and αBC chaperone activity against Aβ(1-42) amyloid aggregation. a) Aβ alone (10 μM) or in the presence of sHSP or the indicated O-GlcNAcylated sHPS protein (1 μM for HPS27 and αBC or 2 μM for αAC) was subjected to aggregation conditions (agitation at 37 °C in a plate reader). Every 5 min, ThT fluorescence (λ_ex_ = 450 nm, λ_em_ = 482 nm) was measured. The y-axis shows fold change in fluorescence compared with the same conditions at t = 0 h. Onset-times were obtained by measuring the time required for fluorescence to reach 3-times the initial reading. Onset-time results are mean ±SEM of four experimental replicates. Statistical significance was determined using a one-way ANOVA test followed by Tukey’s test (Aβ plus HSP27 versus Aβ plus O-GlcNAcylated proteins). b) O-GlcNAcylation activates HSP27 chaperone activity in a substoichiometric fashion. Aβ alone (10 μM) or in the presence of HSP27 or the indicated ratios of HSP27/HSP27(gT184) (2 μM) was subjected to aggregation conditions (agitation at 37 °C in a plate reader). Every 5 min, ThT fluorescence (λ_ex_ = 450 nm, λ_em_ = 482 nm) was measured. The y-axis shows fold change in fluorescence compared with the same conditions at t = 0 h. Onset-times were obtained by measuring the time required for fluorescence to reach 3-times the initial reading. Onset-time results are mean ±SEM of four experimental replicates. Statistical significance was determined using a one-way ANOVA test followed by Tukey’s test.

Next, we tested whether O-GlcNAcylation could improve the activity of sHSPs when it is present at substoichiometric levels in the chaperone oligomers. Accordingly, we incubated different ratios of unmodified HSP27 and HSP27(gT184) for 1 h at 37 °C, which results in subunit exchange and the formation of mixed oligomers^60^with 0, 25, 50, 75, 100% O-GlcNAcylation. We then initiated separate aggregation reactions with 10 μM Aβ and 2 μM of the different ratios of O-GlcNAcylated HSP27 and measured amyloid formation by ThT fluorescence (Figure 4b). Strikingly, we found that as little as 25% HSP27(gT184) in the mixed oligomer was able to significantly increase the onset time of amyloid formation and observed a fairly linear correlation between the amounts of O-GlcNAcylation and the delay in Aβ amyloid formation. These data demonstrate that the increased chaperone activity induced by this modification is not confined to α-synuclein, but is instead a general anti-amyloid feature, and that it can act substoichiometrically.

### O-GlcNAcylation disrupts the ACD-IXI interaction and increases the size of HSP27 oligomers

Next, we set out to test the molecular mechanisms behind O-GlcNAc activation of HSP27 by first examining whether this modification inhibits the binding of the IXI sequence to the chaperone cleft of the ACD domain. In the case of the unmodified IXI sequence, previous experiments showed that the intermolecular interaction between the HSP27’s native IXI peptide (^178^EITIPVTFE^186^) and the ACD domain was too weak to measure reliably. ^40^ However, introduction of a Phe to His mutation at position 185, giving ^178^EITIPVTHE^186^, overcame this limitation. ^40^ Accordingly, we synthesized N-terminally biotinylated peptides corresponding to this improved sequence or the glycopeptide with an O-GlcNAc at Thr184. We then individually immobilized these peptides on a streptavidin-coded microfluidic chip and used surface plasmon resonance (SPR) to measure the binding of monomeric HSP27 ACD domain to these surfaces (Figure 5a). In the case of the unmodified peptide, we observed a K_D_ of 3.14 ± 0.63 μM. In contrast, we detected no binding of the ACD domain to the O-GlcNAcylated peptide. In fact, we saw a negative binding response, which we attribute to a non-specific interaction between the ACD domain and streptavidin that was blocked by the glycopeptide, as we observed in the raw data (Supplementary Figure 4). As confirmation, we synthesized the same two peptides without biotin and measured their binding to the ACD domain by isothermal titration calorimetry (ITC) (Figure 5b). Using this technique, we found a K_D_ of 14.3 ± 1.2 μM for the unmodified peptide and again essentially no binding to the O-GlcNAcylated variant. Importantly, our measured binding constants for the unmodified peptide are consistent with previously published values. ^40^

**Figure 5.**
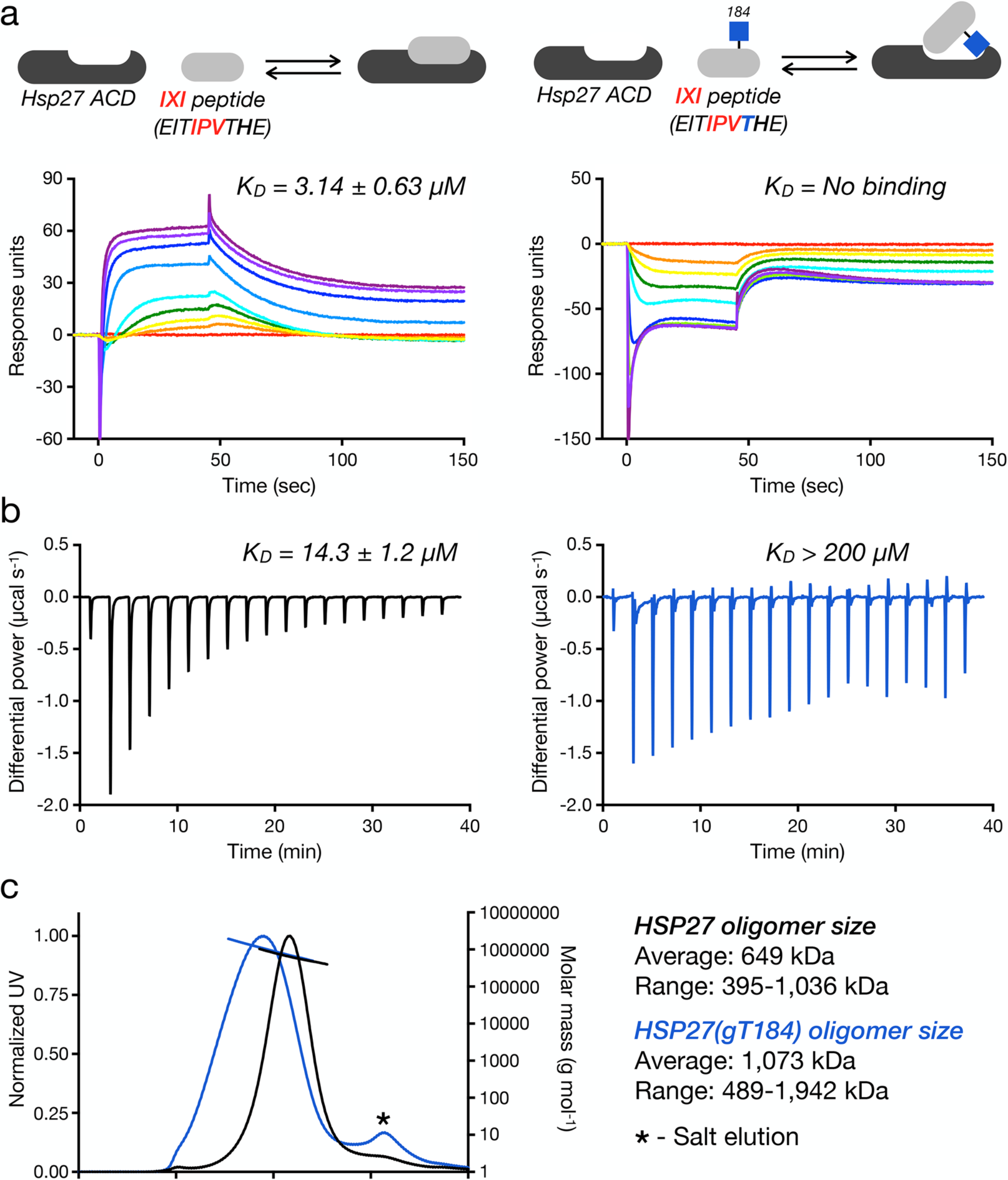
O-GlcNAcylation blocks the IXI/ACD HSP27 domain-interaction and increases the size of HSP27 oligomers. a) The indicated biotinylated IXI or O-GlcNAcylated-IXI peptides were immobilized on a streptavidin-coated SPR chip and the binding of the recombinant HSP27 ACD was measured using surface plasmon resonance (SPR) and the binding constants were determined using Biacore T100 analysis software. b) The corresponding non-biotinylated peptide were titrated against the HSP27 ACD and the binding constant was determined using MicroCal PEAQ-ITC analysis software. c) HSP27 or HSP27(gT184) were analyzed by SEC-MALS showing a larger size and distribution of HSP(gT184) oligomers compared to unmodified HSP27.

The IXI-ACD interaction controls both the accessibility of the chaperone cleft and the oligomer size of HSP27. Therefore, we next used size exclusion chromatography linked to multiple angle light scattering (SEC-MALS) to measure any consequences of O-GlcNAcylation at T184 on HSP27 oligomer size. We found that HSP27(gT184) forms larger oligomers than the unmodified protein (Figure 5c). Specifically, HSP27(gT184) had an average oligomer size of ∼47 monomers, while the unmodified oligomer consisted of only ∼28 monomers. Additionally, the distribution of the HSP(gT184) oligomer size was larger than that of the unmodified protein. Again, the size of our unmodified oligomers are in excellent agreement with previously published data. ^40^ Together, these results are consistent our original hypothesis that O-GlcNAcylation of the IXI domain inhibits its interaction with the ACD chaperone cleft of sHSPs, presumably generating a dynamic structure that can more readily bind to hydrophobic segments and growing amyloid fibers.

## Discussion

Our results demonstrate for the first time that O-GlcNAcylation activates the anti-amyloid activity of all three of the C-terminal IXI-containing sHSPs. We also demonstrate that mechanistically, this is likely due to a decreased physical interaction between the chaperone cleft of the ACD with the IXI-containing C-terminus. We believe that these results have important implications for targeting O-GlcNAc in neurodegenerative diseases. For example, potent inhibitors of OGA are being tested clinically; however, the focus in these and other pre-clinical studies has largely been on the O-GlcNAcylated proteins (e.g., tau) that directly form toxic amyloids. Given the striking substoichiometric activity of sHSPs, and our demonstration that only a fraction of the sHSP must be O-GlcNAcylated for improved activity, we speculate that increasing the modification status of these proteins has an any equally if not more important role for blocking amyloid formation. Interestingly, we also find that O-GlcNAcylation does not appear to affect the parallel role for sHSPs in the maintenance of folded proteins. Specifically, modified HSP27 does not further stabilize the partially unfolded state of the model client protein citrate synthase (Supplementary Figure 5) ^61^. This result suggests that sHSP O-GlcNAcylation could function selectively to prevent protein aggregation rather than globally upregulating all of the protein’s chaperone functions. In summary, we have discovered a new mechanism that further supports a critical role for O-GlcNAcylation in the prevention and potential treatment of neurodegenerative diseases, with key implications for the evaluation of OGA inhibitors as they progress through clinical development.

## METHODS

Methods and any associated references are available in the online version of the paper.

## ACKNOWLEDGMENTS

This research was supported by the National Institutes of Health (R01GM114537) and the Anton Burg Foundation to M.R.P. and the University of Vienna to C.F.W.B.. N.J.P. and S.P.M. were supported by NIGMS T32GM118289, and A.T.B was supported as an USC Dornsife Chemistry-Biology Interface Trainee. SPR, ITC, and SEC-MALS were performed at the USC Nanobiophysics Core Facility. TEM images were collected at the USC Core Center of Excellence in Nano Imaging. ThT measurements were performed at the USC Bridge Institute.

## AUTHOR CONTRIBUTIONS

A.T.B., P.M.L., S.M., N.J.P., S.P.M., T.T.T, C.F.W.B., and M.R.P. designed experiments and interpreted data. A.T.B., P.M.L., N.J.P., and S.P.M. synthesized and purified proteins. A.T.B and P.M.L. performed amyloid aggregation reactions and associated analyses. A.T.B. and T.T.T. performed SPR analysis. A.T.B performed ITC and SEC-MALs analysis. S.M. performed citrate synthase aggregation. A.T.B. and M.R.P. prepared the manuscript.

## ADDITIONAL INFORMATION

Supplementary information is available. Correspondence and requests for materials should be addressed to M.R.P.

## REFERENCES

1. Zachara, N. E. Critical observations that shaped our understanding of the function(s) of intracellular glycosylation (O-Glc NAc). FEBS Lett 592, 3950–3975 (2018).

2. Bond, M. R. & Hanover, J. A. A little sugar goes a long way: the cell biology of O-GlcNAc. J Cell Biol 208, 869–880 (2015).

3. Yang, X. & Qian, K. Protein O-GlcNAcylation: emerging mechanisms and functions. Nat Rev Mol Cell Biol 18, 452–465 (2017).

4. Vocadlo, D. J. O-GlcNAc processing enzymes: catalytic mechanisms, substrate specificity, and enzyme regulation. Curr Opin Chem Biol 16, 488–497 (2012).

5. Shafi, R. et al. The O-GlcNAc transferase gene resides on the X chromosome and is essential for embryonic stem cell viability and mouse ontogeny. Proc Natl Acad Sci USA 97, 5735–5739 (2000).

6. Sinclair, D. A. R. et al. Drosophila O-GlcNAc transferase (OGT) is encoded by the Polycomb group (PcG) gene, super sex combs (sxc). Proc Natl Acad Sci USA 106, 13427–13432 (2009).

7. Yang, Y. R. et al. O-GlcNAcase is essential for embryonic development and maintenance of genomic stability. Aging Cell 11, 439–448 (2012).

8. Yuzwa, S. A. & Vocadlo, D. J. O-GlcNAc and neurodegeneration: biochemical mechanisms and potential roles in Alzheimer’s disease and beyond. Chem Soc Rev 43, 6839–6858 (2014).

9. Wani, W. Y., Chatham, J. C., Darley-Usmar, V., McMahon, L. L. & Zhang, J. O-GlcNAcylation and neurodegeneration. Brain Res. Bull. 133, 80–87 (2017).

10. O’Donnell, N., Zachara, N. E., Hart, G. W. & Marth, J. D. Ogt-dependent X-chromosome-linked protein glycosylation is a requisite modification in somatic cell function and embryo viability. Mol Cell Biol 24, 1680–1690 (2004).

11. Wang, A. C., Jensen, E. H., Rexach, J. E., Vinters, H. V. & Hsieh-Wilson, L. C. Loss of O-GlcNAc glycosylation in forebrain excitatory neurons induces neurodegeneration. Proc Natl Acad Sci USA 113, 15120–15125 (2016).

12. Liu, F., Iqbal, K., Grundke-Iqbal, I., Hart, G. & Gong, C. O-GlcNAcylation regulates phosphorylation of tau: a mechanism involved in Alzheimer’s disease. Proc Natl Acad Sci USA 101, 10804–10809 (2004).

13. Liu, F. et al. Reduced O-GlcNAcylation links lower brain glucose metabolism and tau pathology in Alzheimer’s disease. Brain 132, 1820–1832 (2009).

14. Aguilar, A. L., Hou, X., Wen, L., Wang, P. G. & Wu, P. A Chemoenzymatic Histology Method for O-GlcNAc Detection. ChemBioChem 18, 2416–2421 (2017).

15. Pinho, T. S., Correia, S. C., Perry, G., Ambrósio, A. F. & Moreira, P. I. Diminished O-GlcNAcylation in Alzheimer’s disease is strongly correlated with mitochondrial anomalies. BBA - Molecular Basis of Disease 1865, 2048–2059 (2019).

16. Yuzwa, S. A. et al. Increasing O-GlcNAc slows neurodegeneration and stabilizes tau against aggregation. Nat Chem Biol 8, 393–399 (2012).

17. Borghgraef, P. et al. Increasing brain protein O-GlcNAc-ylation mitigates breathing defects and mortality of Tau.P301L mice. PLoS ONE 8, e84442 (2013).

18. Graham, D. L. et al. Increased O-GlcNAcylation reduces pathological tau without affecting its normal phosphorylation in a mouse model of tauopathy. Neuropharmacology 79, 307–313 (2014).

19. Hastings, N. B. et al. Inhibition of O-GlcNAcase leads to elevation of O-GlcNAc tau and reduction of tauopathy and cerebrospinal fluid tau in rTg4510 mice. Mol Neurodegener 12, 39–16 (2017).

20. Yuzwa, S. A., Cheung, A. H., Okon, M., McIntosh, L. P. & Vocadlo, D. J. O-GlcNAc modification of tau directly inhibits its aggregation without perturbing the conformational properties of tau monomers. J Mol Biol 426, 1736–1752 (2014).

21. Marotta, N. P. et al. O-GlcNAc modification blocks the aggregation and toxicity of the protein α-synuclein associated with Parkinson’s disease. Nat Chem 7, 913–920 (2015).

22. Lewis, Y. E. et al. O-GlcNAcylation of α-Synuclein at Serine 87 Reduces Aggregation without Affecting Membrane Binding. ACS Chem Biol 12, 1020–1027 (2017).

23. Levine, P. M. et al. α-Synuclein O-GlcNAcylation alters aggregation and toxicity, revealing certain residues as potential inhibitors of Parkinson’s disease. Proc Natl Acad Sci USA 116, 1511–1519 (2019).

24. Bukau, B., Weissman, J. & Horwich, A. Molecular chaperones and protein quality control. Cell 125, 443–451 (2006).

25. Hartl, F. U., Bracher, A. & Hayer-Hartl, M. Molecular chaperones in protein folding and proteostasis. Nature 475, 324–332 (2011).

26. Bakthisaran, R., Tangirala, R. & Rao, C. M. Small heat shock proteins: Role in cellular functions and pathology. BBA - Proteins and Proteomics 1854, 291–319 (2015).

27. Haslbeck, M., Weinkauf, S. & Buchner, J. Small heat shock proteins: Simplicity meets complexity. J Biol Chem 294, 2121–2132 (2019).

28. Kappé, G. et al. The human genome encodes 10 alpha-crystallin-related small heat shock proteins: HspB1-10. Cell Stress and Chaperones 8, 53–61 (2003).

29. Kriehuber, T. et al. Independent evolution of the core domain and its flanking sequences in small heat shock proteins. FASEB J 24, 3633–3642 (2010).

30. Jehle, S. et al. Solid-state NMR and SAXS studies provide a structural basis for the activation of alphaB-crystallin oligomers. Nat Struct Mol Biol 17, 1037–1042 (2010).

31. Baldwin, A. J. et al. Quaternary dynamics of αB-crystallin as a direct consequence of localised tertiary fluctuations in the C-terminus. J Mol Biol 413, 310–320 (2011).

32. McDonald, E. T., Bortolus, M., Koteiche, H. A. & Mchaourab, H. S. Sequence, structure, and dynamic determinants of Hsp27 (HspB1) equilibrium dissociation are encoded by the N-terminal domain. Biochemistry 51, 1257–1268 (2012).

33. Baldwin, A. J. et al. Probing dynamic conformations of the high-molecular-weight αB-crystallin heat shock protein ensemble by NMR spectroscopy. J Am Chem Soc 134, 15343–15350 (2012).

34. Hochberg, G. K. A. et al. The structured core domain of αB-crystallin can prevent amyloid fibrillation and associated toxicity. Proc Natl Acad Sci USA 111, E1562–70 (2014).

35. Kudva, Y. C., Hiddinga, H. J., Butler, P. C., Mueske, C. S. & Eberhardt, N. L. Small heat shock proteins inhibit in vitro A beta(1-42) amyloidogenesis. FEBS Lett 416, 117–121 (1997).

36. Raman, B. et al. AlphaB-crystallin, a small heat-shock protein, prevents the amyloid fibril growth of an amyloid beta-peptide and beta2-microglobulin. Biochem J 392, 573–581 (2005).

37. Mainz, A. et al. The chaperone αB-crystallin uses different interfaces to capture an amorphous and an amyloid client. Nat Struct Mol Biol 22, 898–905 (2015).

38. Cox, D., Selig, E., Griffin, M. D. W., Carver, J. A. & Ecroyd, H. Small Heat-shock Proteins Prevent α-Synuclein Aggregation via Transient Interactions and Their Efficacy Is Affected by the Rate of Aggregation. J Biol Chem 291, 22618–22629 (2016).

39. Cox, D. et al. The small heat shock protein Hsp27 binds α-synuclein fibrils, preventing elongation and cytotoxicity. J Biol Chem 293, 4486–4497 (2018).

40. Freilich, R. et al. Competing protein-protein interactions regulate binding of Hsp27 to its client protein tau. Nat Commun 9, 4563 (2018).

41. Delbecq, S. P., Jehle, S. & Klevit, R. Binding determinants of the small heat shock protein, αB-crystallin: recognition of the ‘IxI’ motif. EMBO J 31, 4587–4594 (2012).

42. Pasta, S. Y., Raman, B., Ramakrishna, T. & Rao, C. M. The IXI/V motif in the C-terminal extension of alpha-crystallins: alternative interactions and oligomeric assemblies. Mol. Vis. 10, 655–662 (2004).

43. Hilton, G. R. et al. C-terminal interactions mediate the quaternary dynamics of αB-crystallin. Philos. Trans. R. Soc. Lond., B, Biol. Sci. 368, 20110405 (2013).

44. Rauch, J. N. et al. BAG3 Is a Modular, Scaffolding Protein that physically Links Heat Shock Protein 70 (Hsp70) to the Small Heat Shock Proteins. J Mol Biol 429, 128–141 (2017).

45. Roquemore, E. P. et al. Vertebrate lens alpha-crystallins are modified by O-linked N-acetylglucosamine. J Biol Chem 267, 555–563 (1992).

46. Roquemore, E. P., Chevrier, M. R., Cotter, R. J. & Hart, G. W. Dynamic O-GlcNAcylation of the small heat shock protein alpha B-crystallin. 35, 3578–3586 (1996).

47. Guo, K. et al. Translocation of HSP27 into liver cancer cell nucleus may be associated with phosphorylation and O-GlcNAc glycosylation. Oncol Rep 28, 494–500 (2012).

48. Rambaruth, N. D., Greenwell, P. & Dwek, M. V. The lectin Helix pomatia agglutinin recognises O-GlcNAc containing glycoproteins in human breast cancer. Glycobiology 22, 839–848 (2012).

49. Wang, S. et al. Quantitative proteomics identifies altered O-GlcNAcylation of structural, synaptic and memory-associated proteins in Alzheimer’s disease. J Pathol 243, 78–88 (2017).

50. Deracinois, B. et al. O-GlcNAcylation site mapping by (azide-alkyne) click chemistry and mass spectrometry following intensive fractionation of skeletal muscle cells proteins. Journal of Proteomics 186, 83–97 (2018).

51. Li, J. et al. An Isotope-Coded Photocleavable Probe for Quantitative Profiling of Protein O-GlcNAcylation. ACS Chem Biol 14, 4–10 (2019).

52. Muir, T. W., Sondhi, D. & Cole, P. A. Expressed protein ligation: a general method for protein engineering. Procn Natl Acad Sci USA 95, 6705–6710 (1998).

53. Evans, T. C., Benner, J. & Xu, M. Q. Semisynthesis of cytotoxic proteins using a modified protein splicing element. Protein Sci 7, 2256–2264 (1998).

54. Dawson, P., Muir, T., Clark-Lewis, I. & Kent, S. Synthesis of proteins by native chemical ligation. Science 266, 776–779 (1994).

55. Matveenko, M., Cichero, E., Fossa, P. & Becker, C. F. W. Impaired Chaperone Activity of Human Heat Shock Protein Hsp27 Site-Specifically Modified with Argpyrimidine. Angew Chem Int Ed Engl 55, 11397–11402 (2016).

56. Alderson, T. R. et al. Local unfolding of the HSP27 monomer regulates chaperone activity. Nat Commun 1–16 (2019). doi: 10.1038/s41467-019-08557-8

57. Blanco-Canosa, J. B. & Dawson, P. E. An efficient Fmoc-SPPS approach for the generation of thioester peptide precursors for use in native chemical ligation. Angew Chem Int Ed Engl 47, 6851–6855 (2008).

58. Metanis, N., Keinan, E. & Dawson, P. E. Traceless Ligation of Cysteine Peptides using Selective Deselenization. Angew Chem Int Ed Engl (2010). doi: 10.1002/anie.201001900

59. Shang, S., Tan, Z., Dong, S. & Danishefsky, S. J. An advance in proline ligation. J Am Chem Soc 133, 10784–10786 (2011).

60. Aquilina, J. A., Shrestha, S., Morris, A. M. & Ecroyd, H. Structural and functional aspects of hetero-oligomers formed by the small heat shock proteins αB-crystallin and HSP27. J Biol Chem 288, 13602–13609 (2013).

61. Buchner, J., Grallert, H. & Jakob, U. Analysis of chaperone function using citrate synthase as nonnative substrate protein. Meth Enzymol 290, 323–338 (1998).

