## Supplementary Information for "O-GlcNAcylation of small heat shock proteins enhances their anti-amyloid chaperone activity"

### Table of contents:

**Supplementary Figure 1.** Synthesis and characterization of O-GlcNAcylated HSP27. **Page S2**
**Supplementary Figure 2.** Synthesis and characterization of O-GlcNAcylated αAC. **Page S3**
**Supplementary Figure 3.** Synthesis and characterization of O-GlcNAcylated αBC. **Page S3**
**Supplementary Figure 4.** O-GlcNAcylated IXI-peptide blocks the background ACD-streptavidin interaction in the SPR analysis. **Page S4**
**Supplementary Figure 5.** O-GlcNAcylation does not improve the chaperone activity of HSP27 against citrate synthase aggregation. **Page S4**

**Experimental Methods** **Page S5**

**Supplementary References** **Page S9**

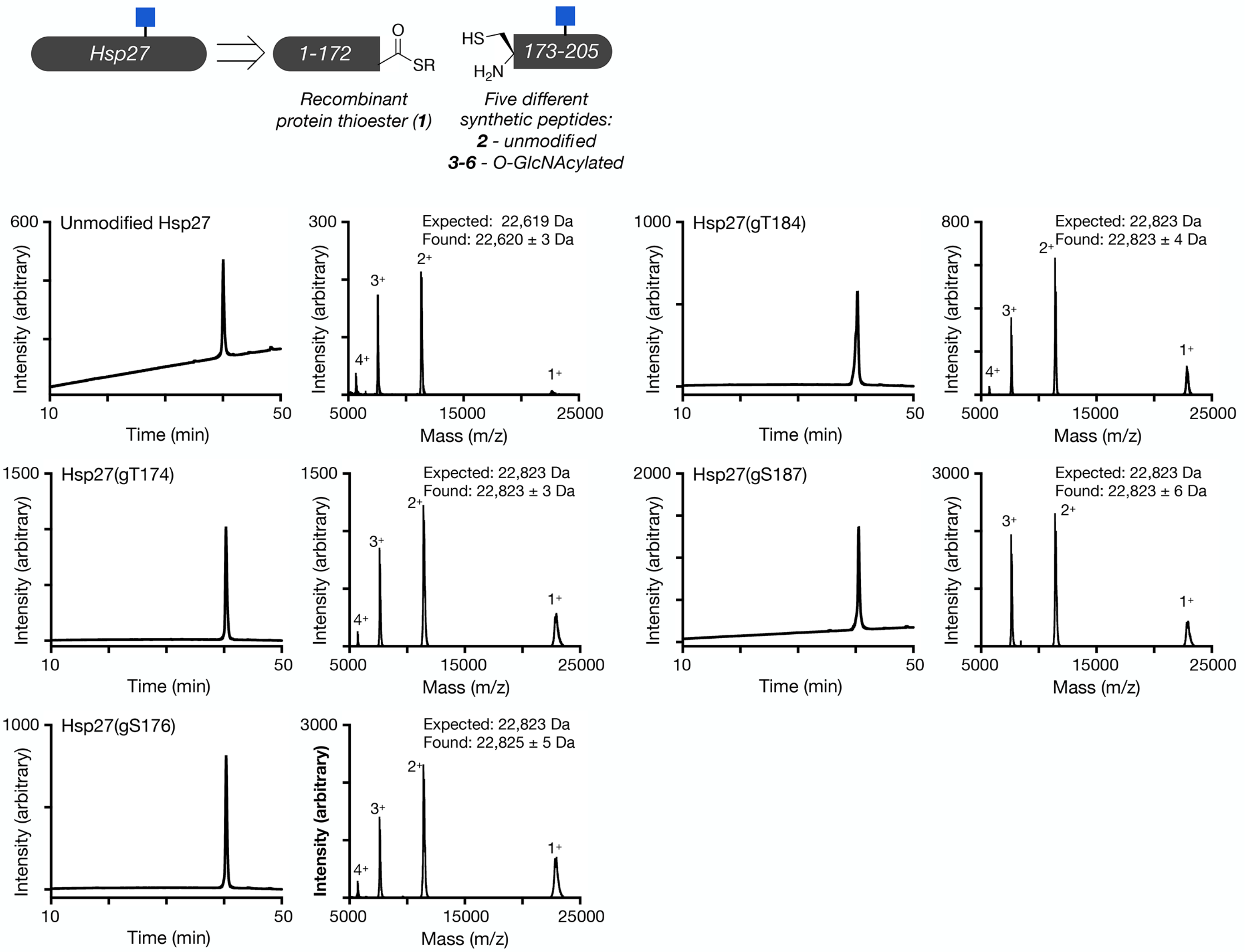


**Supplementary Figure 1. Synthesis and characterization of O-GlcNAcylated HSP27.** Unmodified and differentially O-GlcNAcylated versions of HSP27 were retrosynthetically deconstructed into a recombinant protein thioester and peptides prepared by solid phase peptide synthesis. Analytical RP-HPLC traces and MALDI-TOF-MS of the indicated synthetic proteins.

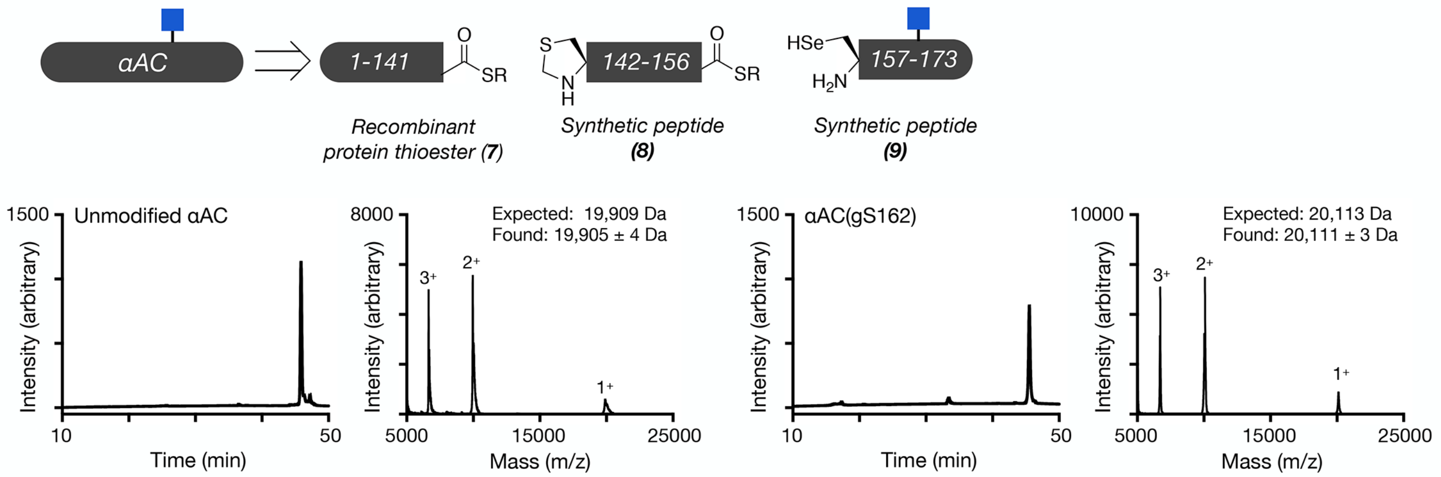


**Supplementary Figure 2. Synthesis and characterization of O-GlcNAcylated αAC**. O-GlcNAcylated αAC was retrosynthetically deconstructed into a recombinant protein thioester and two peptides prepared by solid phase peptide synthesis. Analytical RP-HPLC traces and MALDI-TOF-MS of the indicated recombinant or synthetic proteins.

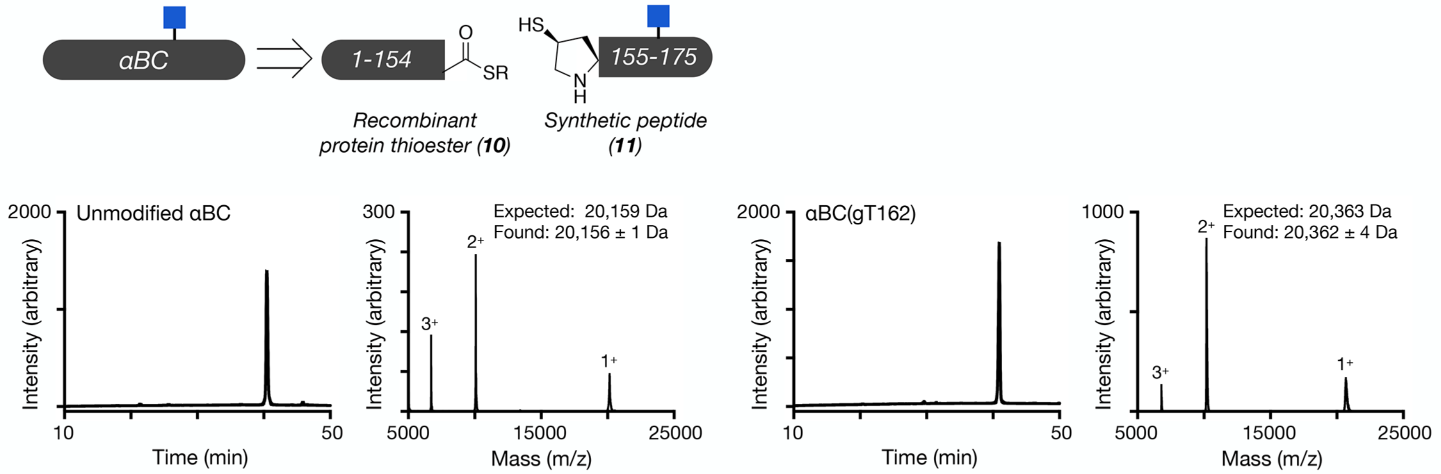


**Supplementary Figure 3. Synthesis and characterization of O-GlcNAcylated αBC**. O-GlcNAcylated αBC was retrosynthetically deconstructed into a recombinant protein thioester and a peptide prepared by solid phase peptide synthesis. Analytical RP-HPLC traces and MALDI-TOF-MS of the indicated recombinant or synthetic proteins.

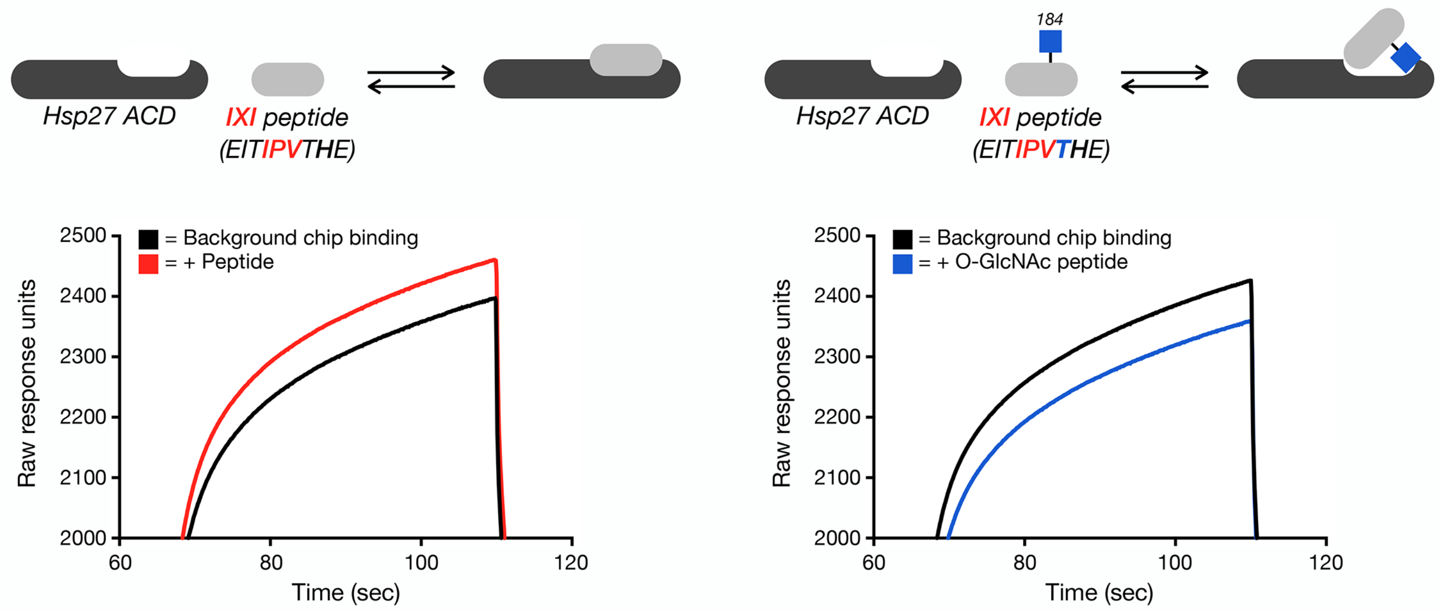


**Supplementary Figure 4. O-GlcNAcylated IXI-peptide blocks the background ACD-streptavidin interaction in the SPR analysis.** Raw SPR data from the experiment in Figure 5a shows less ACD binding to microfluidic chips loaded with the biotinylated, O-GlcNAcylated peptide than background streptavidin alone.

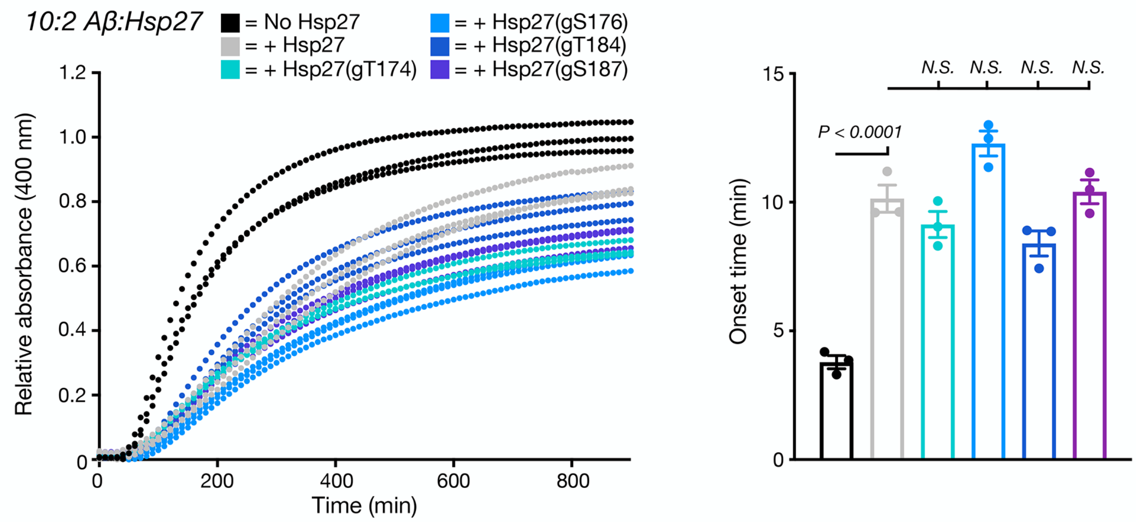


**Supplementary Figure 5. O-GlcNAcylation does not improve the chaperone activity of HSP27 against citrate synthase aggregation.** Citrate synthase (2 µM) in the presence or absence of the indicated HSP27 proteins (0.45 µM) were incubated at 45 °C while measuring the absorbance at 400 nm. Onset-times were obtained by measuring the time required for absorbance to reach 3-times the initial reading. Onset-time results are mean ±SEM of three experimental replicates. Statistical significance was determined using a one-way ANOVA test followed by Tukey's test.

### EXPERIMENTAL METHODS

**General.** All solvents and reagents were purchased from commercial sources and used without any further purification. All aqueous solutions were prepared using ultrapure laboratory grade water (deionized, filtered, and sterilized) obtained from an in-house ELGA water purification system. Growth media were prepared, sterilized, stored, and used according to the instructions of the manufacturer. Antibiotics were prepared as stock solutions at a concentration of 1000× (100 mg/mL ampicillin sodium salt) and stored at -20 °C. All bacterial growth media and cultures were handled using sterile conditions under an open flame. Protein concentrations were determined by the Pierce BCA Protein Assay Kit (Thermo Fisher Scientific). Reversed-phase high-performance liquid chromatography (RP-HPLC) was performed using an Agilent Technologies 1200 Series HPLC instrument with a diode array detector with semi-preparative and analytical C4 or C8 columns obtained from Higgins Analytical. Large scale reversed phase liquid chromatographic purifications were performed on a Biotage Isolera One equipped with SNAP Bio C4 or C18 10g reversed-phased cartridges. The following reversed-phase chromatography buffers were used: buffer A, 0.1% TFA in $H_2O$; buffer B, 0.1% TFA and 90% ACN in $H_2O$. Mass spectra were acquired on an API 3000 LC/MS-MS system (Applied Biosystems/MDS SCIEX) or a Daltonics Autoflex MALDI-TOF (Bruker) using α-cyano-4-hydroxycinnamic acid (HCCA) as the matrix.

**Peptide synthesis.** Peptides were synthesized using standard Fmoc solid-phase chemistry. Unless otherwise stated, pre-loaded Wang resins were used as solid supports. Typical coupling reactions utilize Fmoc-protected amino acids (5 eq.), HBTU (5 eq.) and DIEA (10 eq.) in DMF incubated for 1 h. For glycosylated serine or threonine residues, 2 eq. of Pfp-activated monomer, prepared as previously described[1], in 3 mL of DMF were coupled overnight. Selenocysteine and $\gamma$-thioproline residues were coupled overnight using 2 eq. of Pfp-activated versions of commercially available para-methoxybenzyl- fmoc-selenocysteine or (*2S*,*4R*)-fmoc-mercaptopyrrolidine amino acids (both from ChemImpex). Upon completion of peptide syntheses, acetyl groups were deprotected with hydrazine monohydrate (80% v/v in MeOH) twice for 45 min with mixing. Peptides were then globally deprotected and cleaved (95:2.5:2.5 TFA/$H_2O$/triisopropylsilane) for 4 h at room temperature. For selenocysteine containing peptides, 1.3 eq. of dithiobis(5-nitropyridine) (DTNP) was included in the cleavage cocktail. Cleaved peptides were precipitated out of cold ether, purified by reverse-phase chromatography. and characterized by ESI-MS. Purity was assessed by analytical HPLC. Purified peptides were lyophilized and stored at -20 °C until use.

**Generation of expression plasmids.** HSP27 and αAC expression plasmids were obtained from Addgene (pEGFP-hsp27 wt FL #17444; pE-SUMO-CRYAA #80756) while αBC plasmid (pET28-HSPB5) was a gift from the Benesch lab at the University of Oxford. Desired regions were cloned out using overhang PCR to generate inserts with 5' NdeI and 3' Bpu10I cut sites using KOD HotStart Master Mix (EMD Millipore). After digestion with restriction enzymes (NEB), these were ligated with T4 DNA Ligase (NEB) onto a pTXB1 expression vector at the N-terminus of an AvaE intein bearing a C-terminal 6xHis tag[2]. Ligation mixtures were transformed onto high efficiency DH5α (NEB). After antibiotic selection, plasmids from clones were amplified, purified (QIAGEN Miniprep kit) and analyzed by restriction enzyme digestion. Sequences were confirmed by Sanger sequencing (Laragen) using T7 primers.

**Protein expression and purification.** BL21(DE3) chemically competent *Escherichia coli* (EMD Millipore) cells were transformed with intein-fusion plasmid DNA by heat shock and plated on selective LB agar plates containing 100 µg/mL ampicillin. Single colonies were then inoculated and grown to an $OD_{600}$ of 0.6-0.7 at 37 °C while being shaken at 250 rpm. Expression was induced with IPTG at a final concentration of 1 mM at 37 °C with shaking at 250 rpm for 6 h. After harvesting at 6000g, pellets were resuspended in lysis buffer (50 mM $NaH_2PO_4$, 300 mM NaCl, 1 mM TCEP, 5 mM imidazole, 6 M GuHCl, and 2 mM PMSF, pH 7.5), tip sonicated, and clarified by centrifugation (7000g for 30 min at 4 °C). Protein lysate was loaded onto Co-NTA agarose beads (Genessee Scientific) and washed multiple extensively (50 mM $NaH_2PO_4$, 300 mM NaCl, 2 mM TCEP, 20 mM imidazole, 4 M Urea, pH 7.5). Following washes, protein was eluted (50 mM $NaH_2PO_4$, 300 mM NaCl, 1 mM TCEP, 250 mM imidazole, 4 M Urea, pH 7.5) and dialyzed into PBS to remove excess salts. For protein thioester generation (HSP27 fragment **1**, αAC fragment **7**, αBC fragment **10**), sodium mercaptoethanesulfonate (MESNa) was added to a final concentration of 200 mM and pH was adjusted to 7 prior to overnight incubation at room temperature. For hydrolysis to generate full-length unmodified αAC, αBC, or the HSP27 ACD, dithiothreitol was added to a final concentration of 200 mM and the pH was adjusted to 8 before incubating at 37 °C over 48 h. Thiolysis or hydrolysis reactions were purified by reversed phase liquid chromatography and pure proteins were characterized by analytical RP-HPLC and mass spectrometry. Purified proteins were then freeze-dried yielding lyophilized powders.

Recombinant α-synuclein was expressed from a pRK172 construct containing human wild-type sequence. Expression cultures were grown to an $OD_{600}$ of 0.6-0.7 prior to induction with 0.5 mM final IPTG concentration for 16 h at room temperature. After harvesting, pellets were flash frozen in liquid nitrogen then thawed in a 37 °C bacterial incubator for 20 min. After two more cycles of freeze-thaw, pellets were resuspended in lysis buffer (500 mM NaCl, 100 mM Tris, 10 mM β-mercaptoethanol, 1 mM EDTA, pH 8) and then boiled at 80 °C for 10 min. The bacterial slurry was then allowed to cool down to room temperature after which time PMSF was added to a final concentration of 2 mM. The slurry was mixed by vortexing followed by cooling on ice for 30 min. Protein lysate was clarified by centrifugation. The supernatant was then acidified slowly to pH 3.5 with 1 M HCl, and the solution was incubated on ice to allow proteins to precipitate out. This suspension was centrifuged, and the resulting supernatant was dialyzed overnight against 1% acetic acid in degassed ultrapure water. The dialyzed protein solution was clarified by centrifugation and subjected to RP-HPLC purification, followed by freeze-drying. α-synuclein was purified to >95% purity by analytical HPLC and characterized by ESI-MS.

**Expressed protein ligation to generate HSP27.** Each of the peptide fragments **2-6** (1.1 eq) were dissolved in ligation buffer (300 mM $NaH_2PO_4$, 6 M guanidine HCl, 100 mM MESNa, and 1 mM TCEP, pH 7.4) with 1 eq. of the N-terminal recombinant thioester **1** (3 mM final concentration) and allowed to react at 37 °C until complete as determined by HPLC. Following completion, the reaction was diluted 4-fold into desulfurization buffer (200 mM $NaH_2PO_4$, 3 M guanidine HCl, and 300 mM TCEP, pH 7.0) containing 2% (v/v) ethanethiol, 10% (v/v) tert-butyl-thiol, and the radical initiator VA-061 (as a 0.2 M stock in MeOH). The reaction mixture was stirred at 37 °C for 15 h and then purified by RP-HPLC to yield synthetic unmodified or glycosylated HSP27 with C137A mutation. Proteins were analyzed by analytical HPLC for purity, and masses were confirmed by MALDI-TOF MS. Purified HSP27 variants were lyophilized and stored at -20 °C until use.

**Expressed protein ligation to generate αAC.** Peptide fragment **9** was synthesized on Wang resin using standard protocols above. Peptide fragment **10** was synthesized on Dawson resin (Sigma Aldrich) using standard methods, with the final amino acid coupled using Boc-protected thiazolidine monomer (Advanced Chemtech). After completion of peptide synthesis, the Dawson resin was activated by incubating with 7.5 eq. of para-nitrophenylchloroformate for 1.5 h, followed by 10 eq. of DIEA for 30 min. After peptide cleavage, thiolysis to generate fragment **10** was performed by incubation of the crude peptide mixture in 3M guanidine HCl, 300 mM $NaH_2PO_4$, 200 mM MESNa, pH 7 for 1 h before purification. Lyophilized fragments **9** (1.1 eq) and **10** (1 eq, 7 mM final concentration) were resuspended in ligation buffer (100 mM $NaH_2PO_4$, 6 M guanidine HCl, 100 mM L-ascorbic acid, 250 mM MPAA, and 25 mM TCEP, pH 7). After 16 h at room temperature, product was detected by analytical HPLC and ESI-MS (calculated: 3627.5, found by ESI-MS: 3627.6) when an aliquot was fully reduced with excess TCEP. The entire reaction mixture was purified by preparative RP-LC (Biotage) and the product was isolated as a combination of diselenides or MPAA adducts. The ligation product was then resuspended in deselenization buffer (3M guanidine HCl, 150 mM $NaH_2PO_4$, 25 mM dithiothreitol, 100 m TCEP, pH 5) and incubated for 4 h at room temperature, after which methoxylamine hydrochloride was added to a final concentration of 150 mM to open the thiazolidine ring into a reactive cysteine. After 16 h incubation, the reaction was again purified by reversed phase chromatography. Finally, this intermediate (2 eq.) was dissolved with the N-terminal recombinant thioester fragment **7** (1 eq, 2 mM final concentration) in ligation buffer (300 mM $NaH_2PO_4$, 6 M guanidine HCl, 250 mM MPAA, and 25 mM TCEP, pH 7). The reaction was mixed overnight at room temperature, and product was detected after 16 h. O-GlcNAc S162 αAC was purified by RP-HPLC followed by freeze-drying and storage as lyophilized powder. Purity was assessed by analytical HPLC and mass was characterized by MALDI-TOF MS.

**Expressed protein ligation to generate αBC.** Fragment **10** (1 eq, 3 mM final concentration) and fragment **11** (3 eq) were dissolved in ligation buffer (300 mM $NaH_2PO_4$, 6 M guanidine HCl, 250 mM MPAA, and 25 mM TCEP, pH 7) and allowed to react overnight at room temperature. Product was purified by reversed phase chromatography and lyophilized. This was then resuspended in desulfurization buffer (200 mM $NaH_2PO_4$, 3 M guanidine HCl, and 300 mM TCEP, pH 7.0) at a final concentration of 0.5 mg/mL and allowed to react at 37 °C overnight. Product was purified by RP-HPLC and lyophilized to a powder. Purified O-GlcNAc T162 αBC was characterized by analytical HPLC and MALDI-TOF MS.

**Refolding.** For HSP27 variants, lyophilized proteins were resuspended in 40 mM HEPES·KOH (pH 7.5) at 25 °C for 2 h. For αAC and αBC, lyophilized proteins were resuspended in 6M guanidine-HCl at a concentration of 0.5 mg/mL and dialyzed overnight against 10 mM phosphate buffer, pH 7.4 at 4 °C. Refolded proteins were concentrated and exchanged onto assay buffers using 3K MWCO Amicon-Ultra 0.5 spin filters (Millipore-Sigma).

**α-Synuclein aggregation assays.** Lyophilized recombinant α-synuclein was dissolved with bath sonication in reaction buffer (PBS supplemented with 0.05% sodium azide, pH 7.4). This solution was centrifuged at 20000g for 20 min at 4°C to remove any pre-formed aggregates, and the supernatant was transferred into a fresh tube. Following refolding, sHSPs were buffer exchanged into reaction buffer and concentrated. Protein concentrations of α-synuclein and the sHSPs were determined by BCA assay. Master mixes were prepared by combining the proteins at the final assay concentrations (50 µM α-synuclein and 1 µM sHSP) before dividing into separate 150 µL reaction replicates. The samples were incubated at 37 °C with constant agitation (1000 rpm) in a Thermomixer F1.5 (Eppendorf) for 7 days. At each indicated time point, aliquots were obtained and stored at -80 °C for ThT analysis. In these assay conditions, control aggregation reactions with 1 µM sHSPs (any of the variants) without α-synuclein resulted in no measurable ThT fluorescence increase over the duration of the experiment. Samples from the aggregation assay reaction mixture were diluted in a 96-well plate to a concentration of 1.25 µM with reaction buffer (10 mM PBS, pH 7.4 and 0.05% $NaN_3$) containing 10 µM Thioflavin T (dissolved from a 2000x stock prepared in DMSO). Fluorescence was measured using a Synergy H4 hybrid reader (BioTek). The plate was shaken on "fast" setting for 3 min, followed by data collection ($\lambda_{ex}$ = 450 nm, 9 nm band path, $\lambda_{em}$ = 482 nm, 9 nm band path, reading from the bottom of a plate, gain = 100, read height = 5.00 mm). Fluorescence readings at each time point were normalized to initial fluorescence of pre-aggregation monomers.

**Transmission electron microscopy of aggregation reactions.** For imaging of protein aggregates, a 10 µL (25 uM concentration) droplet from each aggregation experiment was deposited on formvar coated copper grid (150 mesh, Electron Microscopy Sciences) and allowed to sit for 5 min. The excess liquid was removed with filter paper. Grids were then negatively stained for 2 min with 1% uranyl acetate, washed three times with 1% uranyl acetate, each time removing excess liquid with filter paper. The grids were dried for 24 h and then imaged using a JEOL JEM-2100F transmission electron microscope operated at 200 kV, 60,000x magnification, and an Orius Pre-GIF CCD.

**Proteinase K digestion.** Ten micrograms of protein from aggregation reactions were incubated with Proteinase K (Sigma Aldrich P2308) at the indicated concentrations for 30 min at 37 °C. Reactions were quenched by the addition of sample loading buffer (2% final SDS concentration) and boiling at 95 °C for 10 min. Digestion products were separated by SDS-PAGE using precast 12% Bis-Tris gels (Bio-Rad, Criterion XT) with MES running buffer (Bio-Rad). Bands were visualized with Coommassie Brilliant Blue (Bio-Rad).

**Amyloid beta aggregation assays.** Aβ(1-42) (Anaspec) was resuspended in 1% $NH_4OH$ in PBS at a concentration of 10 mg/mL and sonicated to dissolve. This solution was then diluted to 0.5 mg/mL in PBS, pH 7.4 before aliquoting and storing at -80 °C. During each assay, frozen Aβ was slowly thawed on ice before removal of pre-formed aggregates with centrifugation at 20,000g for 30 min at 4 °C. Supernatant was diluted to 10 µM in PBS pH 7.4 with 10 µM ThT dye. This Aβ monomer mix was then divided onto pre-plated sHSPs (previously buffer exchanged into the same assay buffer) at the indicated concentrations. For the mixed HSP27 assay, mixtures of unmodified and gT184 HSP27 at the indicated ratios were pre-incubated at 37 °C for 1 h before pre-plating to facilitate subunit exchange. Using a Synergy H4 Hybrid Plate reader, the microplate was kept at 37 °C and shaken on "fast" setting for 2 min at an interval of 5 min. Reaction was monitored by reading the fluorescence every 5 min over 16 h using the following parameters: ($\lambda_{ex}$ = 450 nm, 9 nm band path, $\lambda_{em}$ = 482 nm, 9 nm band path, reading from the bottom of a plate, gain = 75, read height = 5.00 mm). No measurable increase in fluorescence was measured for any of the sHSPs when incubated without Aβ monomers. Onset was determined using the BioTek Gen5 software by obtaining the time required to reach three times the initial fluorescence reading.

**Surface plasmon resonance.** Peptides were synthesized on Wang resin using standard protocols. N-terminal biotinylation was performed by coupling 1.5 eq. of biotin-PEG4-NHS ester (Click Chemistry Tools) with 0.5 eq. HOBt in DMF overnight. Peptides were purified to >99% purity by analytical HPLC and characterized by ESI-MS (unmodified IXI peptide- calculated: 1511.8 Da, found: 1510.8 Da; O-GlcNAc Thr184 IXI peptide- calculated: 1714.9 Da, found: 1715.2 Da). HSP27 alpha crystallin domain (ACD) was purified from hydrolysis of the intein fusion to >99% purity and characterized by ESI-MS (calculated: 10,751.0 Da, found: 10,751.3 Da).

Experiments were performed on a Biacore T100 system using 10 mM HEPES, 250 mM NaCl, 3 mM EDTA, 0.05% Tween-20, pH 7.4 (HBST) supplemented with 1 mg/mL bovine serum albumin (BSA) as the running buffer. Using 100 nM peptide solutions, 20 response units (RU) of either unmodified or gT184 IXI peptide were immobilized onto the active channels 2 and 4 respectively of a Series S Sensor SA streptavidin chip. Channels 1 and 3 served as reference channels for unmodified or gT184 IXI peptides, respectively. Lyophilized ACD protein resuspended in the running buffer was injected to allow a contact time of 45 sec at a flow rate of 90 µL/min. After 120 sec of dissociation with running buffer, surface was regenerated using a 10 mM glycine, pH 2 buffer for 60s. A concentration course of 125 nM to 120 µM was employed and experiments were repeated three times. Subtracted sensorgrams (active minus reference) were fit using 1:1 binding models to determine affinity using the Biacore T100.

**Isothermal titration calorimetry.** Peptides were synthesized on Wang resin, purified to >99% purity by analytical HPLC and characterized by ESI-MS (unmodified IXI peptide- calculated: 1038.2 Da, found: 1037.4 Da; O-GlcNAc Thr184 IXI peptide- calculated: 1241.2 Da, found: 1240.6 Da). Experiments were performed on a Microcal-PEAQ-ITC system (Malvern). Lyophilized ACD was resuspended in ITC buffer (50 mM $NaH_2PO_4$, 100 mM NaCl, pH 7.4) and was placed in the cell at a concentration of 50 µM. Lyophilized IXI peptides were also resuspended in ITC buffer, adjusted to 1 mM (for unmodified) or 2 mM (for O-GlcNAc Thr184) and loaded onto the titration syringe. Cells were kept at 25 °C and 19 titration injections were applied with 120 sec intervals in between. Only control experiments where peptides were titrated to buffer showed substantial endothermic heats of dilution, hence these control experiments were used to correct peptide-ACD titrations point-by-point. Raw heats were fit using 1:1 binding models to determine affinity of ACD-IXI binding. Experiments were repeated twice.

**Size exclusion chromatography and multiple angle light scattering (SEC-MALS).** Refolded unmodified or gT184 HSP27 proteins (150 µg each) were buffer exchanged to SEC buffer (PBS, pH 7.4) and concentrated to 1.5 mg/mL. Size exclusion was carried out on an Agilent 1200 system equipped with a Shodex 804 column with SEC buffer running at a flow rate of 0.5 mL/min. Molecular weight determination was performed by MALS using a coupled DAWN HELEOS light scattering and rEX refractive index detector (Wyatt Technology Corporation). Data fitting was performed with the ASTRA 6.1 software using a pre-set dn/dc of 0.1850 mL/g.

**Citrate synthase aggregation assays.** The assay was performed according to the literature method with minor modifications. Citrate Synthase (CS) from porcine heart was purchased from Sigma-Aldrich (Taufkirchen, Germany) as an ammonium sulfate suspension then centrifuged to remove most of the ammonium sulfate salts and dialyzed against the storage buffer (50 mM Tris·HCl, 2 mM EDTA, pH 8), final concentration 20-30 µM. The accurate concentration was then determined using bicinchoninic acid (BCA) assay (MW of CS = 48,969 Da) and this stock solution was flash frozen into liquid nitrogen in small aliquots (200-500 µL) and stored at -80 °C. Amorphous aggregation of CS was monitored via measuring the absorbance at 400 nm in a SAFAS UVmc2 double-beam UV-Vis spectrophotometer equipped with a temperature controlled multi-cell holder (SAFAS, Monaco) in 700 µL quartz cuvettes (Hellma Analytics, Germany), 600 µl final volume in triplicate. CS stock solution (obtained as above) was diluted with 40 mM HEPES·KOH (pH 7.5) to a final concentration of 2 µM, and the resulting solution was used as such (control) or treated with Hsp27 variants (0.45 µM final concentration) followed by incubation at 45 °C while measuring the absorbance at 400 nm over 45 min (600 µl final volume in triplicate). Prior to the addition, all Hsp27 variants (lyophilized powders) were freshly dissolved into 40 mM

HEPES·KOH (pH 7.5) buffer, the accurate concentrations of these primary stocks and that of CS were determined via BCA assay and another stock of all Hsp27 samples of concentration 1 mg/mL was prepared to refold them for 3 h at 25 °C. A baseline correction employing only the assay buffer (40 mM HEPES·KOH, pH 7.5) was also performed. The raw data were exported from SAFAS software as Microsoft Excel worksheet and processed using Microsoft Excel and OriginPro. The results were expressed as average relative UV absorbance at 400 nm, where relative absorption at 400 nm = (absorption at 400nm)/(maximal absorption at 400 nm by aggregating CS in the absence of a chaperone).

### REFERENCES


1. De Leon, C. A., Lang, G., Saavedra, M. I. & Pratt, M. R. Simple and Efficient Preparation of O- and S-GlcNAcylated Amino Acids through InBr3-Catalyzed Synthesis of β- N-Acetylglycosides from Commercially Available Reagents. *Org Lett* **20,** 5032–5035 (2018).
2. Shah, N. H., Dann, G. P., Vila-Perelló, M., Liu, Z. & Muir, T. W. Ultrafast Protein Splicing is Common among Cyanobacterial Split Inteins: Implications for Protein Engineering. *J Am Chem Soc* **134,** 11338–11341 (2012).
